# A wealth of novel cell-specific expressed SNVs from tumor and normal scRNA-seq datasets

**DOI:** 10.1101/2022.06.12.495797

**Authors:** Christian Dillard, Evgenia Ulianova, NM Prashant, Hongyu Liu, Nathan Edwards, Anelia Horvath

**Affiliations:** McCormick Genomics and Proteomics Center, School of Medicine and Health Sciences, The George Washington University, Washington, DC, 20037, USA; Departments of Genetics and Genomic Sciences, Icahn School of Medicine at Mount Sinai, New York, NY 10029, USA; Division of Animal Sciences, University of Missouri, Columbia, MO, 65211, USA; Department of Biochemistry and Molecular Medicine, School of Medicine and Health Sciences, The George Washington University, Washington, DC, 20037, USA

**Keywords:** SNV, SNP, mutation, scRNA-seq, single cell, sceSNV, scExecute, somatic mutation, RNA-editing, post-transcriptional modification

## Abstract

We demonstrate a novel variant calling strategy using barcode-stratified alignments on 25 tumor and normal 10XGenomics scRNA-seq datasets (>200,000 cells). Our approach identified 24,528 exonic non-dbSNP single cell expressed (sce)SNVs, a third of which are shared across multiple samples. The novel sceSNVs include unreported somatic and germline variants, as well as RNA-originating variants; some are expressed in up to 17% of the cells, and many are found in known cancer genes. Our findings suggest that there is an unacknowledged repertoire of expressed genetic variants, possibly recurrent and common across samples, in the normal and cancer transcriptome.

## Background

To date, most genetic variation is studied in bulk sequencing datasets, where low (cellular) frequency variants are difficult to distinguish from sequencing errors and other artifacts. Low cellular frequency variants may indicate pre- or early-somatic clonality in cancer and normal tissues or cell-specific RNA post-transcriptional control [1,2]. Furthermore, the ability to detect variants at cell level and knowledge of natural cellular expressed variation are highly compatible with emerging cutting-edge technologies assessing cell-level gene and feature function (Perturb-Seq)[3], characterizing RNA-protein interactions (STAMP), or conducting targeted cell-level genome editing (RADARS) [3–5].

## Results and Discussion

Here, we apply a novel strategy that utilizes barcode-stratified alignments and variant calling on 25 tumor and normal publicly accessible scRNA-seq datasets including prostate cancer (pc), cholangiocarcinoma (chlg), neuroblastoma (nb), normal adrenal (na) and normal embryo (ne (>200,000 cells) generated using the 10XGenomics 3 ‘UTR workflow [6–8]. We extracted single cell alignments using scExecute [9], on pooled scRNA-seq alignments generated by STARsolo [10] and on each single cell alignment called variants using GATK [11] and Strelka2 [12] in parallel, retaining for downstream analysis only variants confidently identified by both callers. The pipeline is presented on Figure 1a.

**Figure 1.**
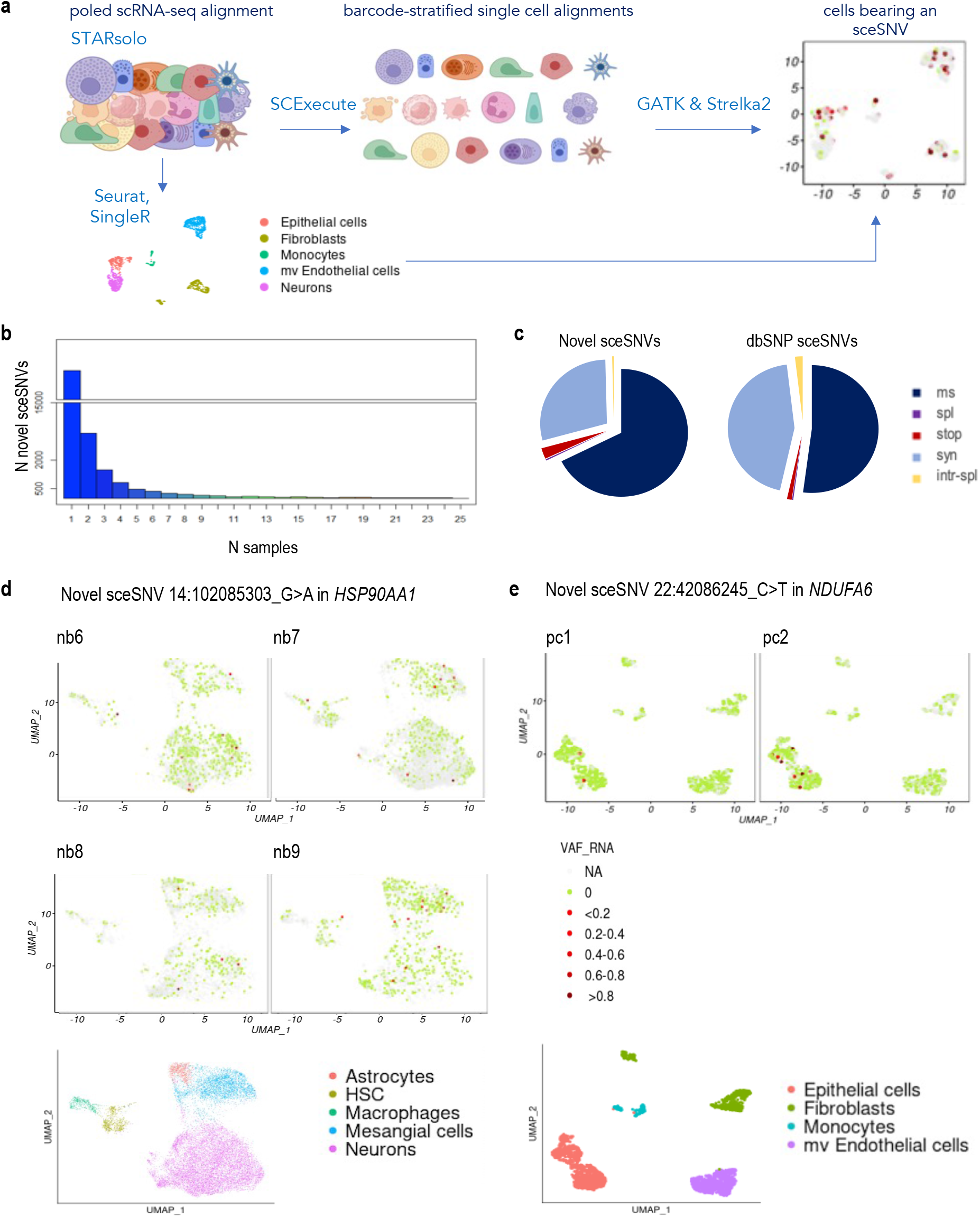
a. Processing pipeline. **b**. Distribution of the number of novel sceSNVs detected in one or more samples. **c**. Distribution of predicted functional annotations between the novel and the previously reported sceSNV; novel sceSNVs are enriched in missense and stop-codon involving substitutions. **d**. top: UMAP projections visualizing the cell distribution and the cellular expressed variant allele frequency (VAF_RNA) of the missense substitution at locus 14:102085303_G>A in the gene *HSP90AA1* across 4 samples from the neuroblastoma dataset. The red color intensity shows the relative expression of the sceSNV in cells with at least 3 sequencing reads covering the sceSNV locus, and the green color indicates that all the reads covering the SNV locus carried the reference nucleotide, consistent with non-zero gene expression. Cells in which the SNV locus is covered by less than 3 reads (corresponding to low or absent gene expression) are shown in grey. Bottom: cell types as classified by SingleR. **e**. top: UMAP projections visualizing the cell distribution of the missense substitution at locus 22:42086245_C>T in the gene *NDUFA6* in the two prostate cancer samples; the expression of the sceSNV is confined to epithelial cells (bottom).

Using this approach, we identified 24,528 exonic non-dbSNP cell-specific expressed single nucleotide variants (sceSNV) from the 25 samples (Table1, and S_Table1), 7,824 of which were observed in more than one sample, and 384 of which are observed in more than half of the datasets (Figure 1b). Of these non-dbSNP sceSNVs, 1,539 (6.3%) are reported in the database of somatic mutations COSMIC [13] and the rest are novel. Some novel sceSNVs were expressed in up to 17% of the cells in a dataset, and many were positioned in known cancer related genes. Cancer genes with multiple novel non-synonymous sceSNVs in more than one sample included *JUN, JAK1, NFKB1A, PIC3R1, RAC1* and *RBX1*. The genes with at least 15 novel non-synonymous sceSNVs in more than one sample were *HSP90AA1, HSP90AB1, ELOB, GSTP* and *JUN*. Novel sceSNVs had a higher proportion of missense and stop-codon involving substitutions, and a lower proportion of synonymous variants, as compared to the dbSNP sceSNVs in the same dataset (p<0.0001, chi-squared test, Figure 1c). In addition, the novel sceSNVs had a higher proportion of A>G substitutions (17.7% vs 14.4% in the DbSNP sceSNVs, S_Figure1). For 1272 of the novel sceSNVs loci, we observed changes of the reference nucleotide into two different alternative nucleotides in different cells and samples, and for 120 loci – changes into all three possible alternative nucleotides (See S_Table1).

**Table 1.**
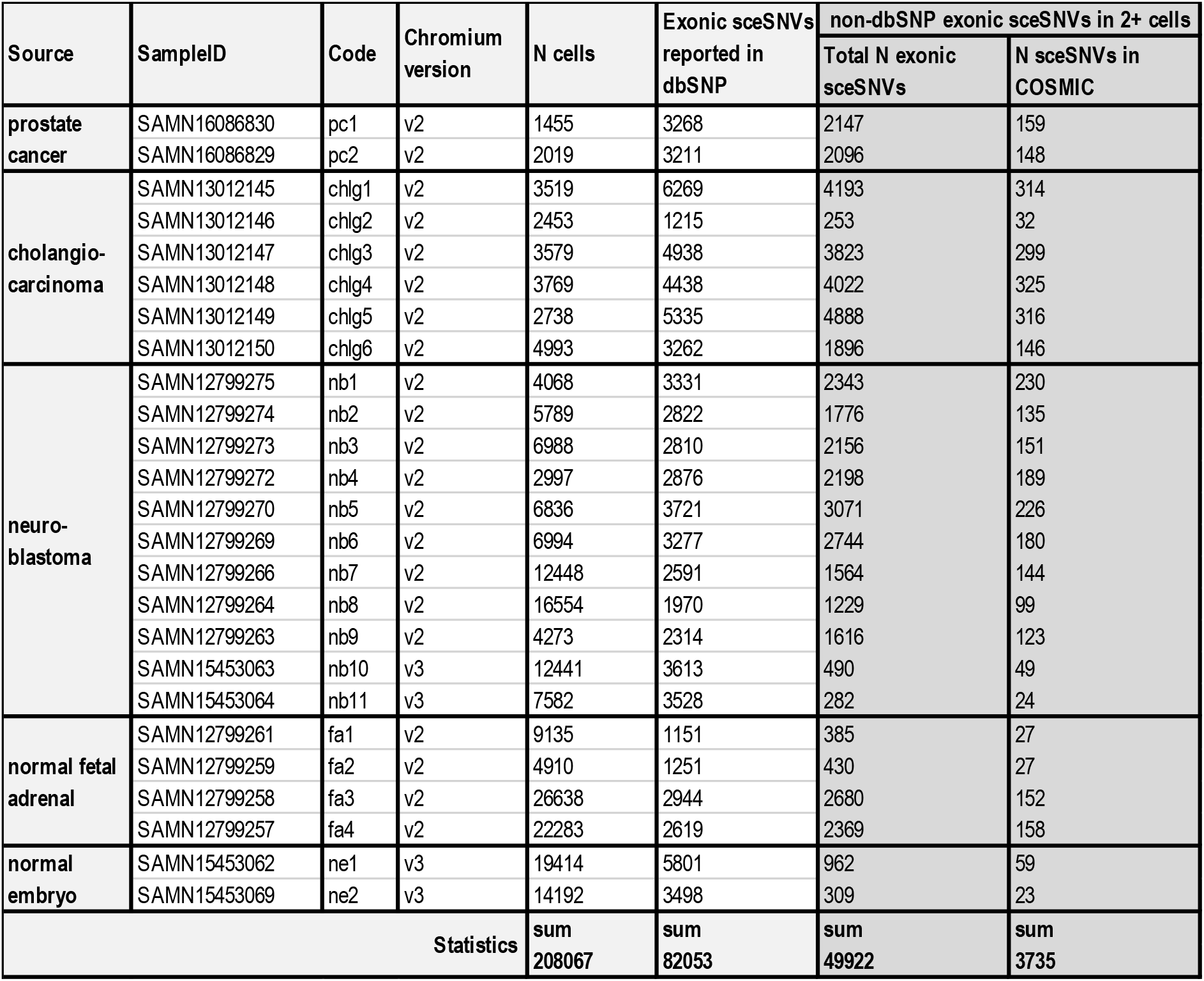
SceSNVs in 2 and more cells across 25 tumor and normal samples

Next, we estimated the expressed Variant Allele Frequency (VAR_RNA, calculated as the proportion of sequencing reads carrying the alternative nucleotide over all the reads covering the locus) [14], and plotted it over the two-dimensional UMAP projection of the cells clustered and annotated based on gene expression [15,16] using scSNVis [17]. While many sceSNVs did not show a distinct cell-distribution pattern (Figure 1d), some were confined to particular cell clusters and cell types despite the ubiquitous expression of their harboring gene (Figure 1e).

Variant calling from individual scRNA-seq alignments is a new and unexplored approach; therefore, to minimize false positives among the novel sceSNVs, we performed stringent quality filtering and examination of the sceSNV confidence at several levels. First, we used for our analyses the intersection of the highest quality calls in at least two cells per dataset by two callers widely used for RNA variant detection, GATK and Strelka2 (S_Methods) [18]. In parallel, for all novel sceSNV positions we estimated the variant read counts across all cells in each dataset using a method for cell-level tabulation of the sequencing read counts bearing reference and variant alleles from barcoded scRNA-seq alignments. [14]. SCReadCounts was fully concordant with the variant call results, identifying variant reads in all cells where sceSNV was called. Third, for 500 arbitrarily selected sceSNVs, we visually reviewed the local alignment using the Integrated Genomics Viewer (IGV) [19]. This analysis showed that for up to 2% of the calls per sample the originally called variant participates in a more complex alteration, involving two or more consecutive nucleotides. These calls were removed across the entire dataset by assessment the presence of more than one call within any 25 consecutive bases in the same cells. The remaining loci showed high quality alignment and confident variant presence (examples on S_Figure2, S_Figure3 and S_Figure4). Fourth, for the same set of 500 sceSNVs we examined the between-cell occurrence and distribution. Many sceSNVs show preferential presence in particular cell-clusters, suggesting a relationship between the sceSNV and cluster-specific gene expression (Figure 1e and S_Figure5) and the related molecular context of cell-specific sceSNV. Fifth, we restricted our analysis to exonic regions, comprising the best-known reference variational context; we additionally filtered out difficult to assess repeated genomic regions and removed ribosomal genes due to their significant number of homologous genes and pseudogenes. We note that while these stringent measures are likely to increase false negatives, in this pilot approach we prioritize high-quality, high confidence sceSNV calls over a comprehensive search for sceSNVs.

The identified novel sceSNVs include previously unreported somatic and germline variants, as well as RNA-originating variation. Without cell-level matched DNA, the origin of sceSNVs is difficult to assign. However, for some sceSNVs, likely origin can be inferred from their cellular allele expression (VAF_RNA) and distribution across cells and samples. For example, while the above-described filtering cannot completely exclude germline DNA variants, they are unlikely to constitute a large proportion of novel sceSNVs because they are not reported in DbSNP and are observed in only a modest proportion of the cells per sample. On the other hand, low cellular frequency sceSNVs are consistent with both somatic DNA sceSNVs and sceSNVs resulting from post-transcriptional RNA-modifications such as RNA-editing. Somatic sceSNVs observed in multiple samples are likely to be reported in COSMIC, suggesting that the non-COSMIC novel sceSNVs are enriched in post-transcriptional RNA-modifications. While our set of novel sceSNVs do not contain previously reported RNA-editing loci and we exclude repetitive genome regions, known to contain the highest frequency of editing events, low cellular frequency editing of exonic positions is possible [20,21]. Another possibility for some non-COSMIC somatic sceSNVs is that they exist only in the period between their incidence and cell death, for example, if they impair critical mechanisms for cell-survival or replication. Such a scenario would prevent replication of cells bearing the sceSNV and result in low cellular frequency, and consequently challenge detection by bulk sequencing techniques. In regard to VAF_RNA, RNA-editing sceSNVs are likely to have higher VAF_RNA variance and attain any value between 0 and 1, whereas for many somatic mutations in biallelically expressed genes VAF_RNA is expected to be closer to 0.5. Finally, sceSNVs resulting from random transcription errors are unlikely to have high prevalence among our set as they are expected to be seen in a single molecule (i.e., represented by only one read per cell), which calls are excluded by our stringent quality filtering. Of note, regardless of DNA- or RNA-origin, the sceSNVs represent part of the functional cellular transcriptome, can exert effects on the proteins sequence and function, and can increase molecular variation of the cell at the multi-omics level.

Finally, to assess potential links between the novel sceSNVs and cell-specific gene expression, for a subset of sceSNVs we performed differential expression analysis between sceSNV-bearing cells and the rest of the cells in the dataset using Deseq2 [22]. Notably, for many sceSNV, we observed gene expression differences concordant across different samples, and consistent with current knowledge. For example, DE analysis of cells with and without the novel stop codon substitution 1:45511398_C>T in *PRDX1* identified 5 and 7 significantly deregulated genes in the two prostate cancer samples respectively (padj < 0.2), four of which are shared between the two samples and deregulated in the same direction (S_Table2). Three of these four genes – *TAGLN, ACTA2* and *MYL9* - participate in a well-known network (S_Figure6). The observation of shared and concurrently deregulated genes in cells bearing the same sceSNVs provides additional evidence for these sceSNVs as true positives and suggests causative, mechanistic and functional implications. Possible scenarios include sceSNVs regulating the expression of one or a set of genes, or RNA-modification events taking place in cells with similar gene expression, both potentially more frequent for sceSNVs co-clustered in cells of similar types.

## Conclusions

Here, we explore for the first-time expressed genetic variation at cell-level. Our findings suggest that there is an unappreciated repertoire of cell-level expressed genetic variation, possibly recurrent and common across samples, that participates in transcriptome function and dynamics in both cancer and normal cells. While the DNA-or RNA-origin of these variants is currently difficult to confidently determine, their appearance and, for some, relationship to certain gene-sets and cell types, suggests novel mechanisms and function for expressed genetic variation. We also demonstrate an assessment strategy for cellular and functional context by studying deregulated genes in the cells bearing specific sceSNVs. Furthermore, correlation between their VAF_RNA and the expression of harboring or other genes using scReQTL [23] may also provide needed biological context. The analysis of sceSNVs from scRNA-seq data is crucial to support emerging methods that use cell-level introduction and tracking of RNA-variants for manipulating cellular behavior and temporal deconvolution of cellular events [3–5]. Interpretation of these new methods’ results will require prior knowledge of naturally occurring cell-level genetic and transcript variation, which we explore in this work. The study of expressed cell-specific variants in scRNA-seq data, as demonstrated here, has the potential to link expressed variation to tissue evolution and cell fate and has a role in the successful implementation of emerging single-cell biotechnologies.

## Methods

The methods used to perform this study are described in detail in S_Methods.

