## Supplementary material for "A wealth of novel cell-specific expressed SNVs from tumor and normal scRNA-seq datasets": S_Methods

### Supplementary Methods

##### *Sequencing Datasets*

To exemplify scExecute we utilized publicly available scRNA-seq data from three different cancer studies providing prostate cancer tissue [1], neuroblastoma, normal fetal adrenal and normal embryo [2], and cholangiocarcinoma [3]. The experimental design and the data generation are described in detail in the original studies. All the studies utilized 10x Genomics Chromium Single Cell 3′ Workflow protocol for the libraries' generation, and the libraries were sequenced on an Illumina NextSeq 500 platform to a length of 150 nt. The sequencing datasets were downloaded from the NCBI Sequence Read Archive (SRA) (accessed on 1 April 2021) under the accession numbers listed in Table1.

##### *Alignment and Variant call*

The pooled raw scRNA-seq sequencing reads were aligned using the STARsolo module of STAR v.2.7.7a in 2-pass mode, with transcript annotations assembly GRCh38.79 [4]. Transcriptome-wide variant calling was performed on each single cell binary alignment map file (SCbam) using SCExecute in conjunction with the HaplotypeCaller module of GATK v.4.2.0.0 and Strelka2 v.2.9.10 in parallel; both tools were used in their default setting [5,6]. The GATK HaplotypeCaller was preceded by the assignment of read groups using the GATK module AddOrReplaceReadGroups, followed by splitting reads that contain Ns in their cigar string with the GATK module SplitNCigarReads.

Our analysis was confined on SNVs (i.e. indels and other variants were filtered out). The sceSNV calls from the individual alignments were filtered using the bcftools utility (v.1.10.2) of SAMtools, retaining sceSNVs with QUAL (Phred-scaled probability) > 100, MQ (mapping quality) > 60, and QD (quality by depth) > 2. SNV loci were annotated using SeattleSeq v.16.00 (dbSNP build 154), and SNVs positioned in non-repetitive regions and non-ribosomal genes were retained for further analysis. Local alignments were visually reviewed through the Integrative Genomics Viewer (IGV, [7]).

##### *Cell-level gene expression assessments*

To estimate cell-level gene expression, we used read-count matrices with the raw gene counts per cell generated by STARsolo. We normalized and scaled the expression data using the SCTransform function, as implemented in Seurat v.3.0 [8]. The cell-feature distributions were then plotted to identify and filter out the outliers and low-quality cells, which we defined after examination of the cell feature distribution. Specifically, based on the cell and feature distribution, we have filtered out: (1) cells with mitochondrial gene expression of above between 10% and 20%, (2) cells with fewer than 1000 genes, and (3) cells with more than between 3500 and 5500 detected genes (to remove potential doublets). The Seurat-processed gene expression values were also used to remove batch effects and cell cycle effects, and further processed for cell type assessments.

##### *Cell Types Classifications*

To define likely cell types with known cell types, we used SingleR v.1.0.5 [9], as previously described [10–12]. SingleR defines likely cell types, comparing genome-wide expression profile of each cell to a database of reference cells' whole transcriptome expression (BluePrint + ENCODE datasets). To select the expression profile corresponding to known cell type, the analysis is rerun iteratively with the top cell types from the previous step.

##### *$VAF_{RNA}$ Estimation and SNV Distribution Plotting*

Single-cell level VAF_RNA was assessed from the pooled scRNA-seq alignments using scReadCounts v.1.1.4, as we have previously described [12]. When provided with barcoded scRNA-seq alignments and genomic loci of interest (with alleles), SCReadCounts tabulates the reference and variant read counts ($n_{ref}$ and $n_{var}$, respectively), and generates a cell-SNV matrix with the $VAF_{RNA}$ estimated at a user-defined threshold of the minimum number of required sequencing reads (minR) for a confident $VAF_{RNA}$ assessment. For the analysis presented herein, we used minR ≥ 3, which excludes from the estimation those positions covered by an insufficient number of reads (in this case 3). The cell-SNV VAF_RNA matrix is than used as an input together with outputs of Seurat and SingleR to plot the SNV-distributions on the two-dimensional UMAP projections using scSNVis [13].

##### *Differential Gene Expression*

Differential gene expression between cells bearing particular sceSNV of interest and the rest of the cells in the dataset was performed on the generated by Seurat raw gene count matrices using Deseq2 [14]. Correction for multiple trials was performed using the module incorporate in Deseq2, and genes with adjusted p-value below 0.2 were considered significantly deregulated.

##### *Comparative Statistical Assessments*

To compare distribution of functional annotations between the novel sceSNVs and sceSNVs reported in dbSNP we used 2x2 contingency tables and chi-square test; p-values below 0.05 were considered significant.

##### *References*

1. Ma X, Guo J, Liu K, Chen L, Liu D, Dong S, et al. Identification of a distinct luminal subgroup diagnosing and stratifying early stage prostate cancer by tissue-based single-cell RNA sequencing. Mol Cancer. 2020;

2. Dong R, Yang R, Zhan Y, Lai H-D, Ye C-J, Yao X-Y, et al. Single-Cell Characterization of Malignant Phenotypes and Developmental Trajectories of Adrenal Neuroblastoma. Cancer Cell. 2020;

3. Zhang M, Yang H, Wan L, Wang Z, Wang H, Ge C, et al. Single-cell transcriptomic architecture and intercellular crosstalk of human intrahepatic cholangiocarcinoma. J Hepatol. 2020;73:1118–30.

4. Kaminow B, Yunusov D, Dobin A. STARsolo: accurate, fast and versatile mapping/quantification of single-cell and single-nucleus RNA-seq data. bioRxiv. 2021;

5. Van der Auwera GA, Carneiro MO, Hartl C, Poplin R, del Angel G, Levy-Moonshine A, et al. From fastQ data to high-confidence variant calls: The genome analysis toolkit best practices pipeline. Curr Protoc Bioinforma. 2013;

6. Kim S, Scheffler K, Halpern AL, Bekritsky MA, Noh E, Källberg M, et al. Strelka2: fast and accurate calling of germline and somatic variants. Nat Methods. 2018;

7. Robinson JT, Thorvaldsdóttir H, Winckler W, Guttman M, Lander ES, Getz G, et al. Integrative genomics viewer. Nat. Biotechnol. 2011. p. 24–6.

8. Butler A, Hoffman P, Smibert P, Papalexi E, Satija R. Integrating single-cell transcriptomic data across different conditions, technologies, and species. Nat Biotechnol. 2018;

9. Aran D, Looney AP, Liu L, Wu E, Fong V, Hsu A, et al. Reference-based analysis of lung single-cell sequencing reveals a transitional profibrotic macrophage. Nat Immunol. 2019;

10. Liu H, Prashant NM, Spurr LF, Bousounis P, Alomran N, Ibeawuchi H, et al. scReQTL: an approach to correlate SNVs to gene expression from individual scRNA-seq datasets. BMC Genomics [Internet]. 2021;22:40. Available from: https://doi.org/10.1186/s12864-020-07334-y

11. Prashant N, Liu H, Dillard C, Ibeawuchi H, Alsaeedy T, Chan KH, et al. Improved SNV discovery from barcode-stratified scRNA-seq alignments. Genes (Basel). 2021;12.

12. Prashant NM, Alomran N, Chen Y, Liu H, Bousounis P, Movassagh M, et al. SCReadCounts: estimation of cell-level SNVs expression from scRNA-seq data. BMC Genomics [Internet]. 2021 [cited 2021 Sep 23];22:689. Available from: https://bmcgenomics.biomedcentral.com/articles/10.1186/s12864-021-07974-8

13. Hongyu Liu, Prashant NM, Nawaf Alomran, Pavlos Bousounis, Mercedeh Movassagh, Nathan Edwards and AH. scSNVis: cell-level visualization of expressed SNVs from scRNA-seq data.

14. Love MI, Huber W, Anders S. Moderated estimation of fold change and dispersion for RNA-seq data with DESeq2. Genome Biol. 2014;
