## Supplementary material for "A wealth of novel cell-specific expressed SNVs from tumor and normal scRNA-seq datasets": S_Figure1

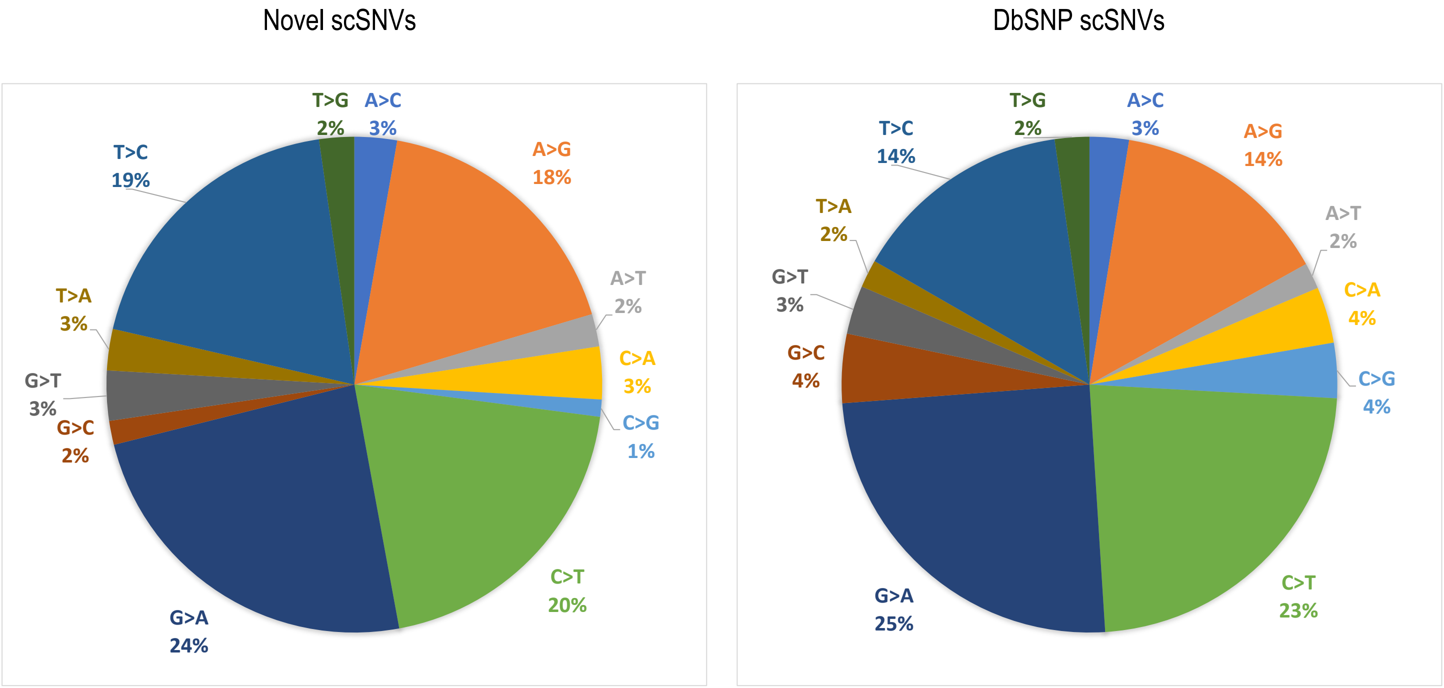


**S_Figure2**. Distribution of predicted functional annotation in novel sceSNVs and sceSNVs overlapping with DbSNP SNVs. The novel sceSNVs contained higher proportions of A>G and T>C substitutions, and lower proportions of C>T and C>G substitutions.
