## Supplementary material for "A wealth of novel cell-specific expressed SNVs from tumor and normal scRNA-seq datasets": S_Figure2

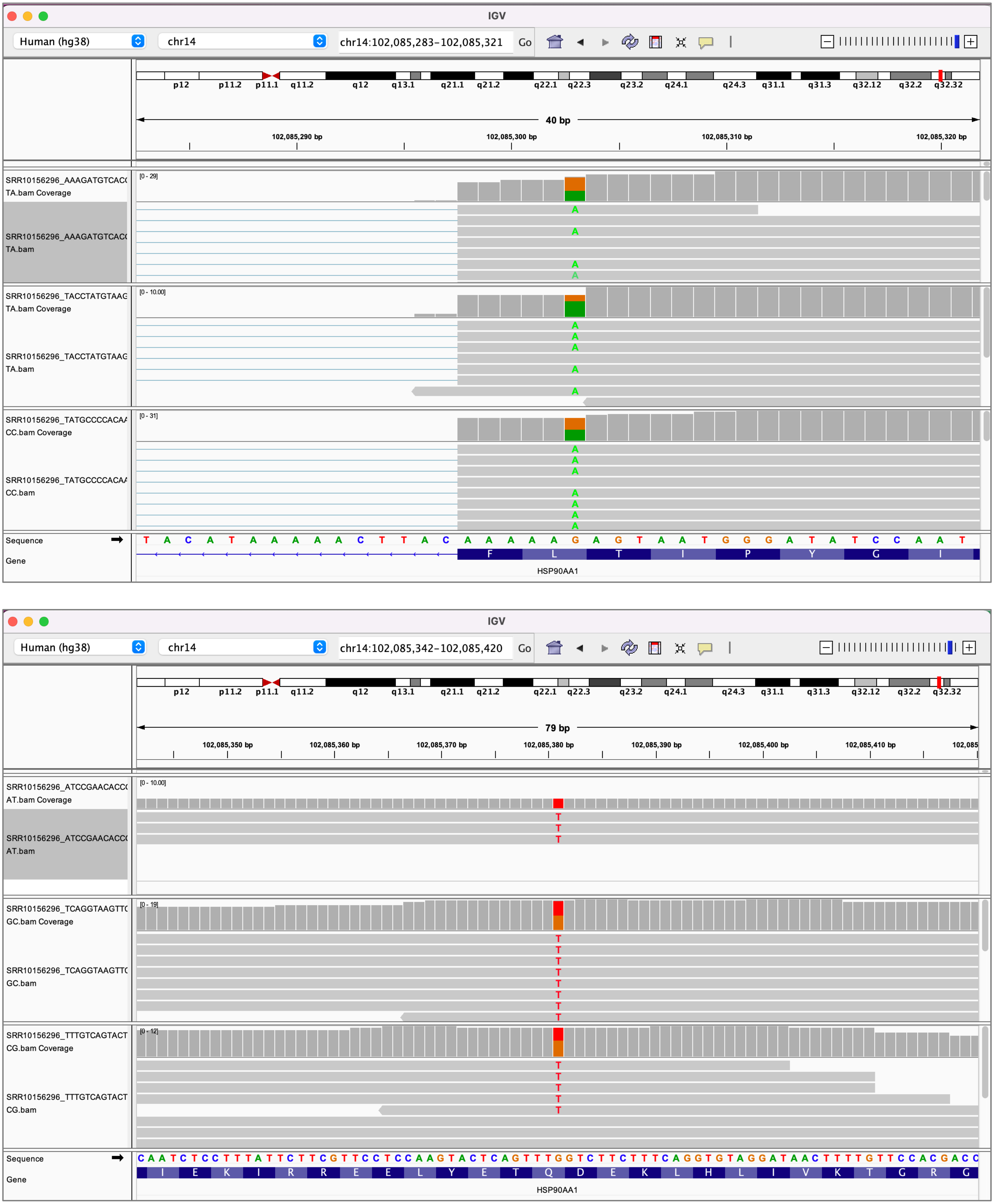


**S_Figure2**. IGV visualization of sceSNV positive cells showing mono-and bi-allelic expression supported by multiple unique sequencing reads of the sceSNV 14:102085303_G>A (top) and the sceSNV in locus 14:102085381_G>T in the gene *HSP90AA1* (bottom).
