## Supplementary material for "A wealth of novel cell-specific expressed SNVs from tumor and normal scRNA-seq datasets": S_Figure3

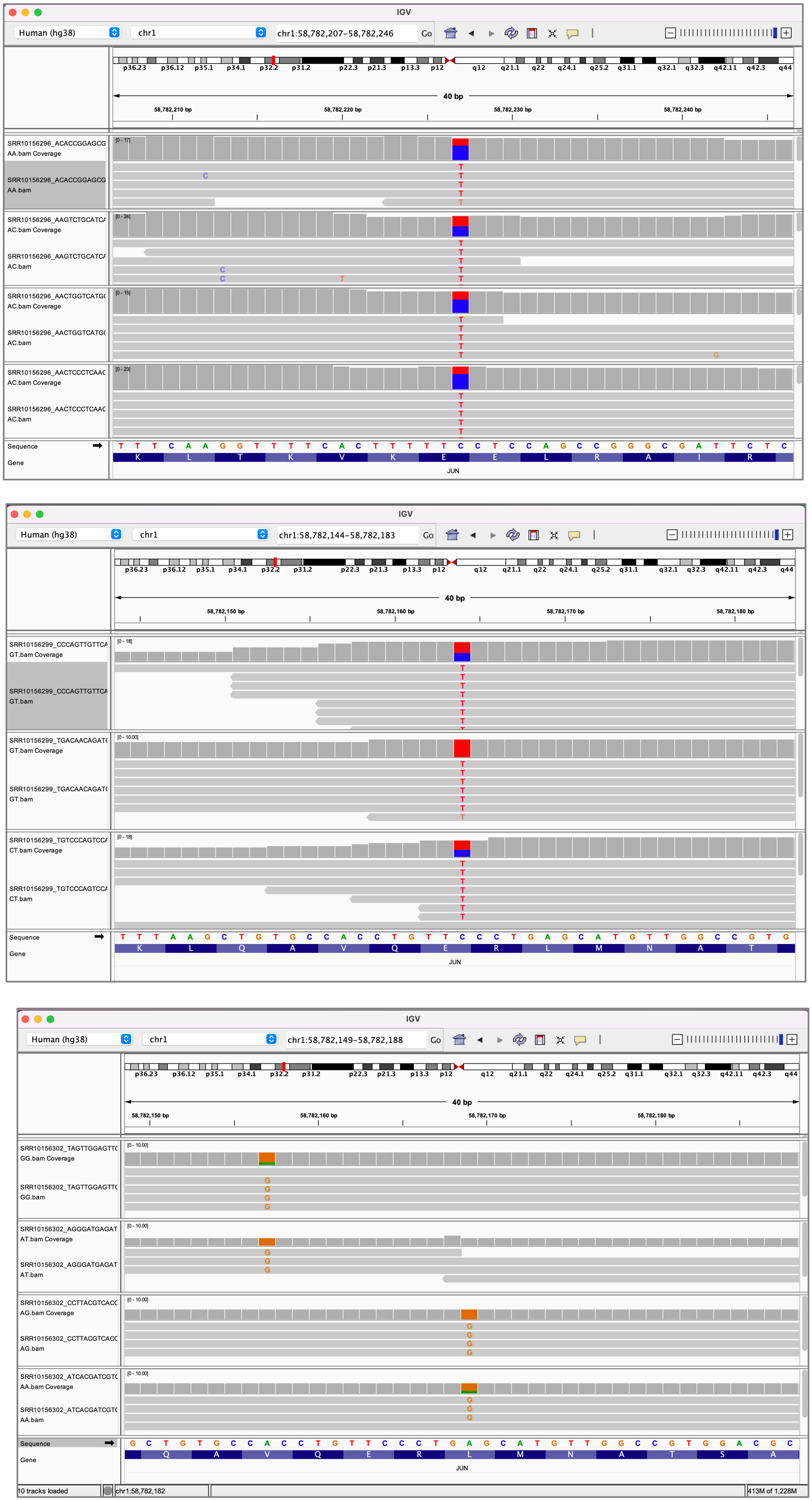


**S_Figure3.** IGV visualization of sceSNV positive cells showing mono-and bi-allelic expression supported by multiple unique sequencing reads of the sceSNV 1:58782227_C>T (top), the sceSNV i1:58782166_C>T (middle), and the sceSNVs 1:58782169_A>G and 1:58782185_A>G (bottom) in the gene *JUN*.
