## Supplementary material for "A wealth of novel cell-specific expressed SNVs from tumor and normal scRNA-seq datasets": S_Figure4

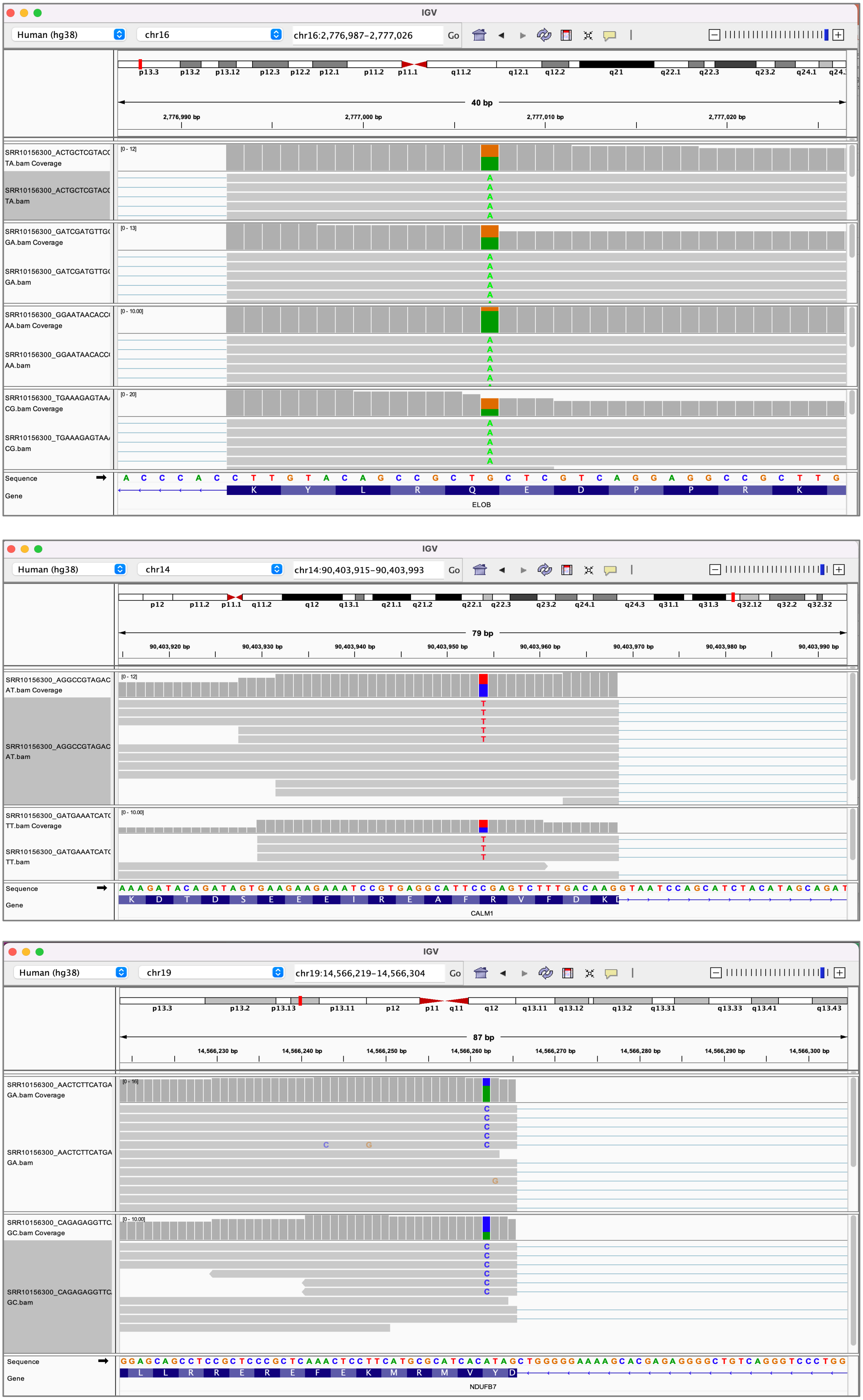


**S_Figure4**. IGV visualization of sceSNV positive cells showing mono-and bi-allelic expression supported by multiple unique sequencing reads of the sceSNV 16:2777007_G>A (top) in the gene *ELOB*, sceSNV 14:90403954_C>T in the gene *CALM1* (middle), and sceSNV 19:14566251_A>C in the gene *NDUFB7*.
