## Supplementary material for "A wealth of novel cell-specific expressed SNVs from tumor and normal scRNA-seq datasets": S_Figure5

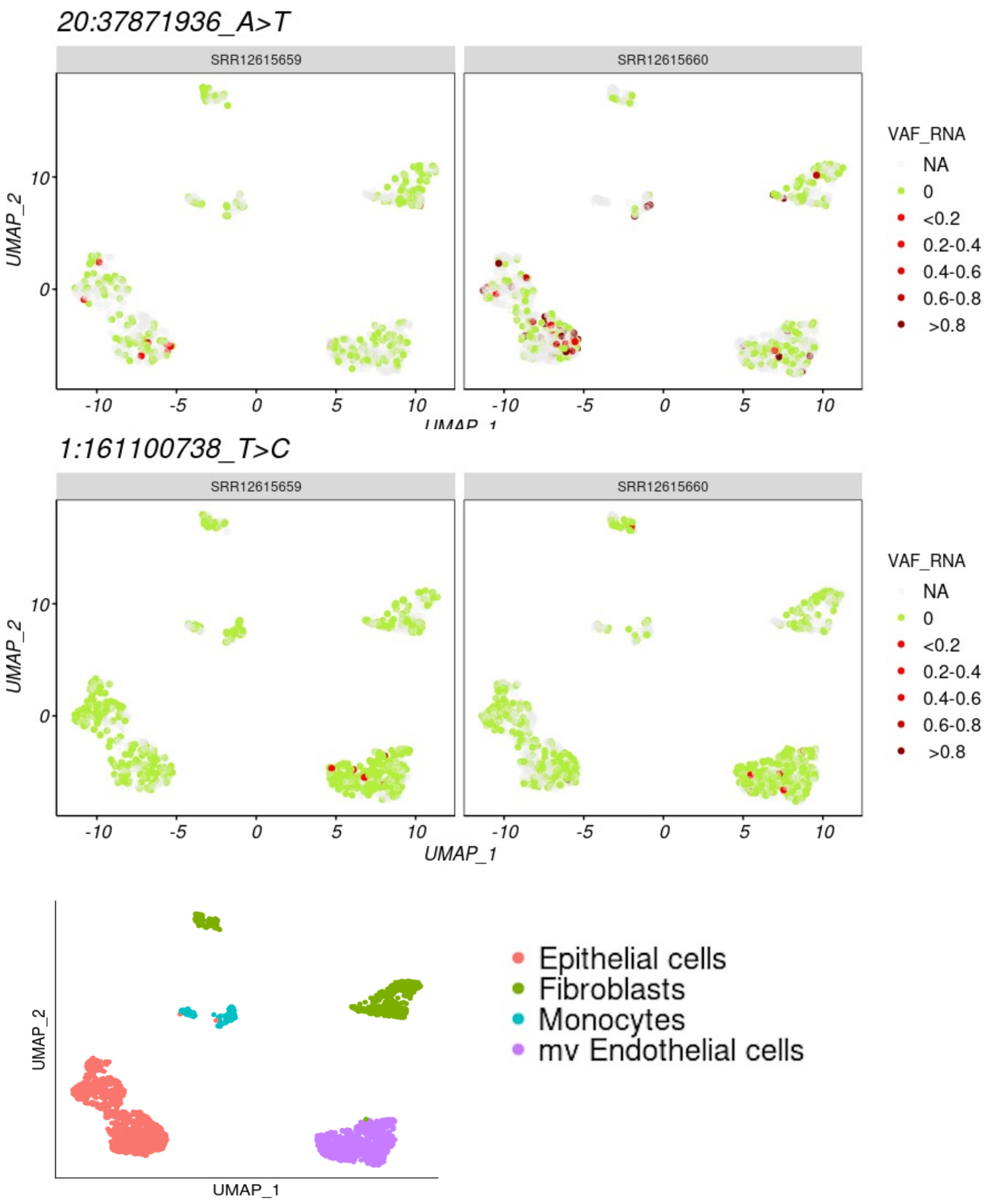


**S_Figure4.** UMAP projections visualizing the cell distribution and the cellular expressed variant allele frequency (VAF_RNA) of the novel sceSNV 20:37871936 _A>T in the gene *CTNNBL1*, seen more frequently in the endothelial cells (top) and sceSNV 1:161100738 in the gene *PFDN2*, which was confined to the endothelial cells. The red color intensity shows the relative expression of the sceSNV in cells with at least 3 sequencing reads covering the sceSNV locus, and the green color indicates that all the reads covering the SNV locus carried the reference nucleotide, consistent with non-zero gene expression. Cells in which the SNV locus is covered by less than 3 reads (corresponding to low or absent gene expression) are shown in grey. Cell types as classified by SingleR are shown at the bottom.
