## Supplementary material for "A wealth of novel cell-specific expressed SNVs from tumor and normal scRNA-seq datasets": S_Figure6

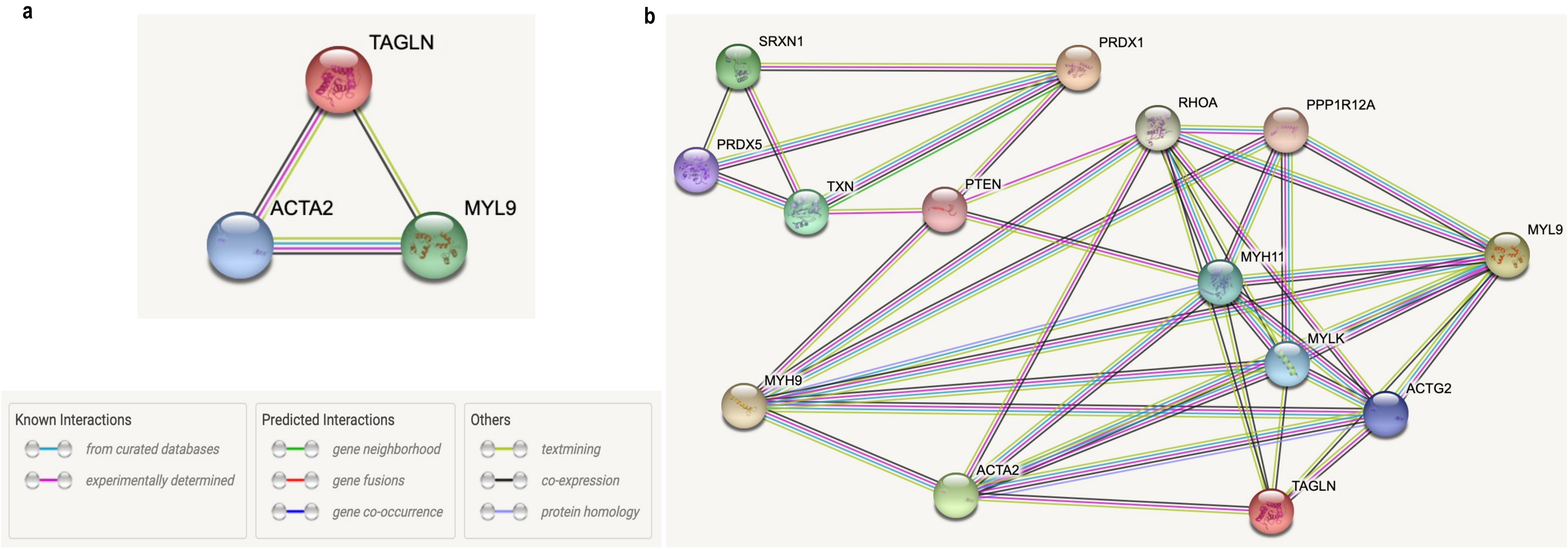


**S_Figure5. a.** Known molecular networks between *ACTA2*, *TAGLN* and *MYL9*, all significantly downregulated in cells expressing the novel stop-codon sceSNV 1:45511398_C>T in the gene *PRDX1*. **b.** Possible links between *PRDX1*, *ACTA2*, *TAGLN* and *MYL9* include *PTEN*, *RHOA* and *MYH11*.
